# Depletion of m^6^A-RNA in *Escherichia coli* reduces the infectious potential of T5 bacteriophage

**DOI:** 10.1101/2024.05.08.593107

**Authors:** Bibakhya Saikia, Sebastian Riquelme-Barrios, Thomas Carell, Sophie Brameyer, Kirsten Jung

## Abstract

*N*^6^-methyladenosine (m^6^A) is the most abundant internal modification of mRNA in eukaryotes that plays, among other mechanisms, an essential role in virus replication. However, the understanding of m^6^A RNA modification in prokaryotes, especially in relation to phage replication, is limited. To address this knowledge gap, we investigated the effects of m^6^A RNA modification on phage replication in two model organisms: *Vibrio campbellii* BAA-1116 (previously *V. harveyi* BB120) and *Escherichia coli* MG1655. An m^6^A-RNA depleted *V. campbellii* mutant (Δ*rlmF*Δ*rlmJ*) did not differ from the wild type in the induction of lysogenic phages or in susceptibility to the lytic Virtus phage. In contrast, the infection potential of the T5 phage, but not that of other T phages or the lambda phage, was reduced in an m^6^A-RNA depleted *E. coli* mutant (Δ*rlmF*Δ*rlmJ*) compared to the wild type. This was shown by a lower efficiency of plaquing and a higher percentage of surviving cells. There were no differences in T5 phage adsorption rate, but the mutant exhibited a 5 min delay in the rise period during the one-step growth curve. This is the first report demonstrating that *E. coli* cells with lower m^6^A RNA levels have a higher chance of surviving T5 phage infection.

**Importance:** The importance of RNA modifications has been thoroughly studied in the context of eukaryotic viral infections. However, their role in bacterial hosts during phage infections is largely unexplored. Our research delves into this gap by investigating the effect of host m^6^A-RNA modifications during phage infection. We found that an *E. coli* mutant depleted of m^6^A-RNA is more resistant to T5 infection than the wild type. This finding emphasizes the need to further investigate how RNA modifications affect the fine-tuned regulation of individual bacterial survival in the presence of phages to ensure population survival.

## Introduction

Chemical modifications of RNA molecules are reversible and dynamic and influence their fate and many cellular processes (1). Of the numerous RNA modifications discovered to date, the eukaryotic m^6^A modification machinery and its presence in host and viral RNA are especially well-studied and provide a potential target for antiviral therapeutics (1–7). However, despite the wealth of knowledge regarding m^6^A-RNA modification in eukaryotes and eukaryotic viruses, the potential implications of m^6^A-RNA modifications in bacteria are limited and in relation to bacteriophages (phages) are still largely unexplored.

Phages are prokaryotic viruses that infect bacteria and use them as hosts to propagate. They are ubiquitous and known to be highly abundant in various natural ecosystems such as soil and water, in addition to the human microbiome (8). Their varied and omnipresent lifecycles reflect their complex interactions with bacterial hosts, which influence their population, virulence and evolution (9). These complex interactions have attracted considerable research interest related to the intricate fine-tuning between hosts and phages. For example, phage-encoded transfer RNAs (tRNAs) are modified like the corresponding host tRNAs, suggesting that they are processed by the same host enzymes (10). Similarly, another study showed that the bacterial toxin CmdT (part of the toxin–antitoxin–chaperone system) acts as an ADP-ribosyltransferase that modifies phage mRNA to block protein translation (11). Thus, as with most eukaryotic viruses, RNA modifications in phages may be performed by host enzymes and may have pro- or anti-viral functions.

Adenosine methylation at the nitrogen (*N*^6^-methyladenosine - m^6^A) is present in all types of *Escherichia coli* RNA (12), however, the methyltransferases are only well documented for ribosomal RNA (rRNA) and tRNA. In *E. coli*, 23S rRNA molecules are methylated at nucleotides A1618 and A2030 by RlmF and RlmJ, respectively (13, 14). In addition, the valine-specific tRNA is methylated at nucleotide A37 by TrmM (YfiC) (15). Notable, RlmF is a structural homolog of METTL16, a mRNA methyltransferase in eukaryotes (16). Overexpression of *rlmF* leads to a slight growth defect in *E. coli* (13), whereas deletion of *rlmJ* leads to a higher growth rate under anaerobic conditions (14). Single deletions of either *rlmF* or *rlmJ* do not interfere with ribosome assembly or influence the growth rate in *E. coli* (17). RlmJ is conserved in the phylum Proteobacteria, whereas RlmF is predominantly found in the class *Gammaproteobacteria* (17). Consequently, RlmF, RlmJ and TrmM are well conserved in the marine bacterium *Vibrio campbellii* BAA-1116 (previously known as *Vibrio harveyi* BB120), a model organism for quorum sensing (18). Our previous work with *V. campbellii* has shown the presence of four intact prophages in the *V. campbellii* genome, two of which can be activated under oxidative (mitomycin C) and heat stress (19).

To explore the potential role of m^6^A-RNA modification in lysogenic and lytic phages, we tested *V. campbellii* and *E. coli* m^6^A-RNA depletion mutants lacking both *rlmF* and *rlmJ,* which encode the major m^6^A rRNA methyltransferases. We started our study by the induction of lysogenic phages in *V. campbellii* and could show that m^6^A modification has no significant effect on the prophage induction. We then searched for lytic phages that showed different infection efficiency in the m^6^A-RNA depleted mutants. We identified bacteriophage T5, whose infectious potential was reduced in the m^6^A-RNA depleted *E. coli* mutant, resulting in higher host survival.

## Results

### m^6^A-RNA depletion in *V. campbellii* does not affect prophage induction or susceptibility to a lytic phage

*V. campbellii* has four intact prophages in its genomes (19). Prophages are viral DNA segments that have integrated into the host genome (also called lysogen) and provide a competitive advantage to the host (20). In *V. campbellii*, ΦHAP-1-like prophage and *Vibrio* kappa-like prophage are triggered into their lytic cycle under oxidative or heat stress (19). Prophage induction triggers a transition from the lysogenic to the lytic cycle, resulting in the activation of dormant viral DNA and subsequent viral replication and host cell lysis. Knowing the effects of oxidative and heat stress on prophage induction in the wild type of *V. campbellii*, we wanted to determine whether the m^6^A-RNA depleted mutant (Δ*rlmF*Δ*rlmJ*) shows a different response. Deletion of *rlmF* and *rlmJ* in *V. campbellii,* which encode the two m^6^A-specific rRNA methyltransferases, resulted, in a decrease of 96.5 % in m^6^A levels of the tRNA-depleted mRNA and rRNA fraction (Fig. 1A).

**FIG 1.**
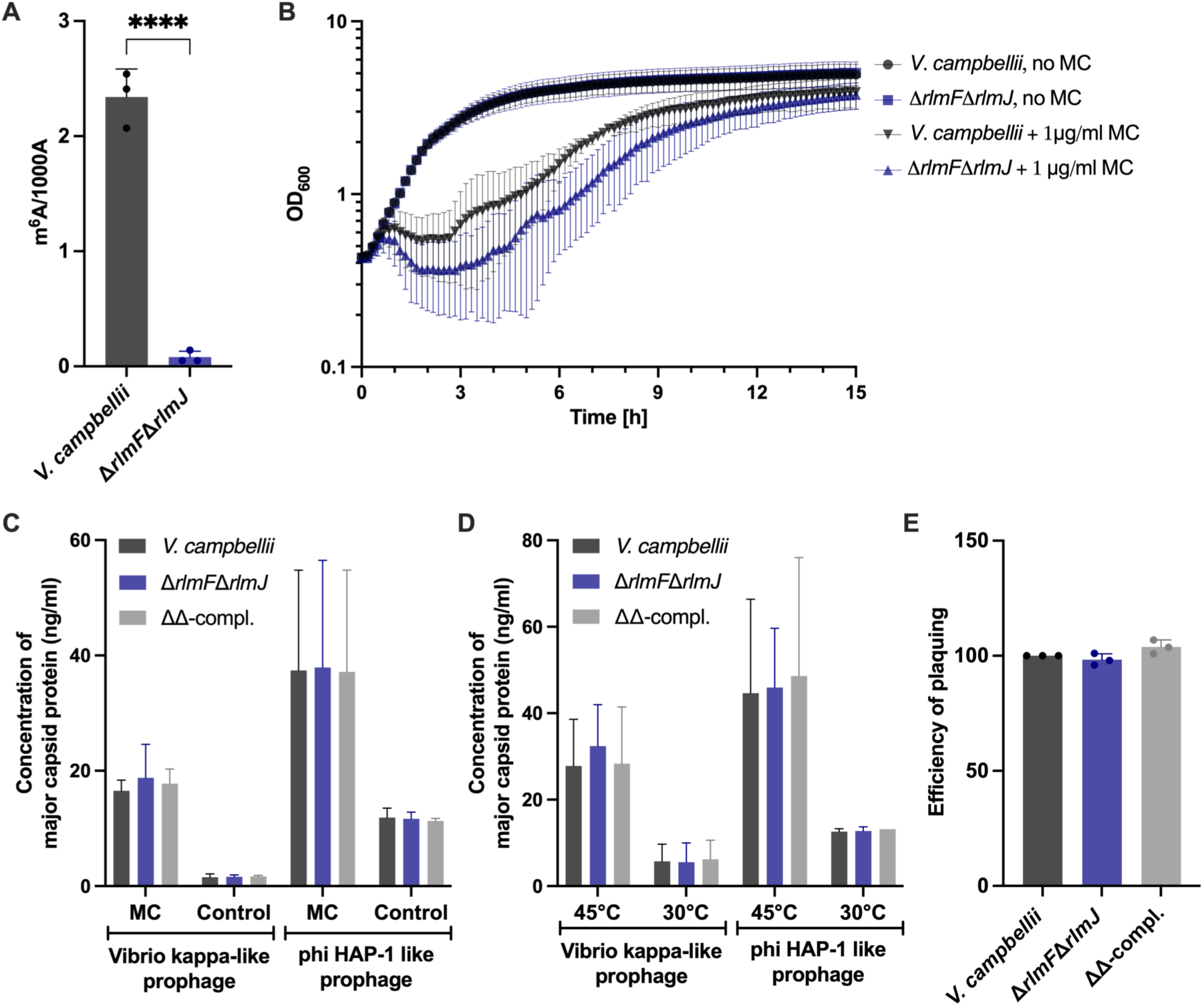
The effect of m^6^A depletion in *V. campbellii* ATCC BAA1116 on the induction of lysogenic phages and susceptibility to lytic Virtus phage. (A) Quantitative analysis of m^6^A related to 1000 A (adenosine) of isolated rRNA+mRNA from *V. campbellii* and its Δ*rlmF*Δ*rlmJ* mutant. (B) The indicated strains were grown to the exponential growth phase and exposed to 1 μg/mL mitomycin C (MC) for 30 minutes. Afterwards, cells were resuspended in fresh medium and optical density (OD_600_) was monitored over time. (C and D) Lysogenic phages were induced in the indicated strains with 1 μg/mL mitomycin C (MC) for 30 minutes (C) or heat at 45°C for 30 minutes (D). Culture supernatants were collected 2 hours after induction and subjected to indirect ELISA to estimate the release of Vibrio kappa-like prophages and ΦHAP-1-like prophages. (E) The indicated strains were grown to the exponential growth phase and infected with phage Virtus. The graph shows the efficiency of plaquing (EOP), which is calculated by dividing the number of plaques obtained from the mutant by the number of plaques obtained from the wild type. The EOP of the wild type was set to 100. Data are presented as the mean ± standard deviation of at least three independent experiments. **** *p* < 0.0001 (Student’s *t*-test).

First, we tested the role of m^6^A-RNA on lysogeny. We exposed the wild type and the Δ*rlmF*Δ*rlmJ* mutant to mitomycin C and monitored the cell lysis caused by phage release over time. At 2.5 hours after exposure to mitomycin C, a greater extent of cell lysis was observed in the Δ*rlmF*Δ*rlmJ* mutant compared to the wild type, but the effect was not statistically significant (Fig. 1B). In parallel, we developed an assay for the quantification of phage particles. For this purpose, we raised antibodies against two major capsid proteins, one for the Vibrio kappa-like prophage and one for the ΦHAP-1-like prophage (VIBHAR_05027 and VIBHAR_01983).

Using an indirect ELISA assay, we found a slightly higher concentration of the capsid protein of the Vibrio kappa-like prophage, indicating a higher release of phage particles from the m⁶A-RNA depleted mutant of *V. campbellii* compared to the wild type after exposure to mitomycin C (Fig. 1C) and heat stress (Fig. 1D). Although this difference was restored in the complemented mutant (ΔΔ-compl.), in which the two genes were chromosomally reintegrated (Fig. 1C and D), the effect was not statistically significant. In the untreated cells, a low level of capsid proteins was observed due to spontaneous induction of prophages (Fig.1C and D).

Under both oxidative and heat stress, we detected higher levels of the capsid protein of the ΦHAP-1-like prophage, indicating a greater release of phages (Fig. 1C and D). However, we could not detect any difference between the wild type, the m^6^A-RNA depleted mutant, or the complemented mutant (Fig. 1C and C).

We then tested whether the m^6^A-RNA modification has an effect during lytic phage infection (Fig. 1E). The availability of the vibriophage Virtus, which is capable of infecting a broad range of *Vibrio* spp. (including *V. campbellii*) (21), provided the unique opportunity to evaluate its infectivity for *V. campbellii* wild type and its Δ*rlmF*Δ*rlmJ* mutant. We tested the efficiency of plaquing by phage Virtus after infection of exponentially grown *V. campbellii* wild type, its Δ*rlmF*Δ*rlmJ* mutant and the complemented mutant (ΔΔ-compl.). However, infectivity by Virtus was comparable for all three strains (Fig. 1E). We also tested the efficiency of plaquing by phage Virtus after infection of stationary phase cells of the *V. campbellii* strains, but could not detect any difference (data not shown).

In summary, the m^6^A-RNA modification had no effect on prophage induction or replication of the lytic Virtus phage in *V. campbellii*.

### m^6^A-RNA depletion in *E. coli* affects bacteriophage T5 infection

Since further exploration of the role of m^6^A RNA modification in *V. campbellii* has been hampered by the lack of easily available lytic phages, we proceeded with *E. coli*, which, although containing cryptic prophages in its genome (22), offers a diverse array of well-characterized lytic phages (23). An m^6^A-RNA depleted mutant (Δ*rlmF*Δ*rlmJ*) *E. coli* MG1655, which has a reduction in the m^6^A-RNA level comparable to that of *V. campbellii* (Fig. 1A), and a complemented mutant were already available (24).

We screened the most prominent lytic phages for *E. coli,* five of the “T-phages” (23, 26) and the bacteriophage λ (27), for their infection potential of the Δ*rlmF*Δ*rlmJ* mutant compared to the wild type (Fig. 2). All phages tested are double-stranded DNA phages belonging to the order *Caudovirales* and characterized by their tail and icosahedral head (28). They represent three families with different specificity for bacterial host receptors (29) (Fig. 2).

**FIG 2.**
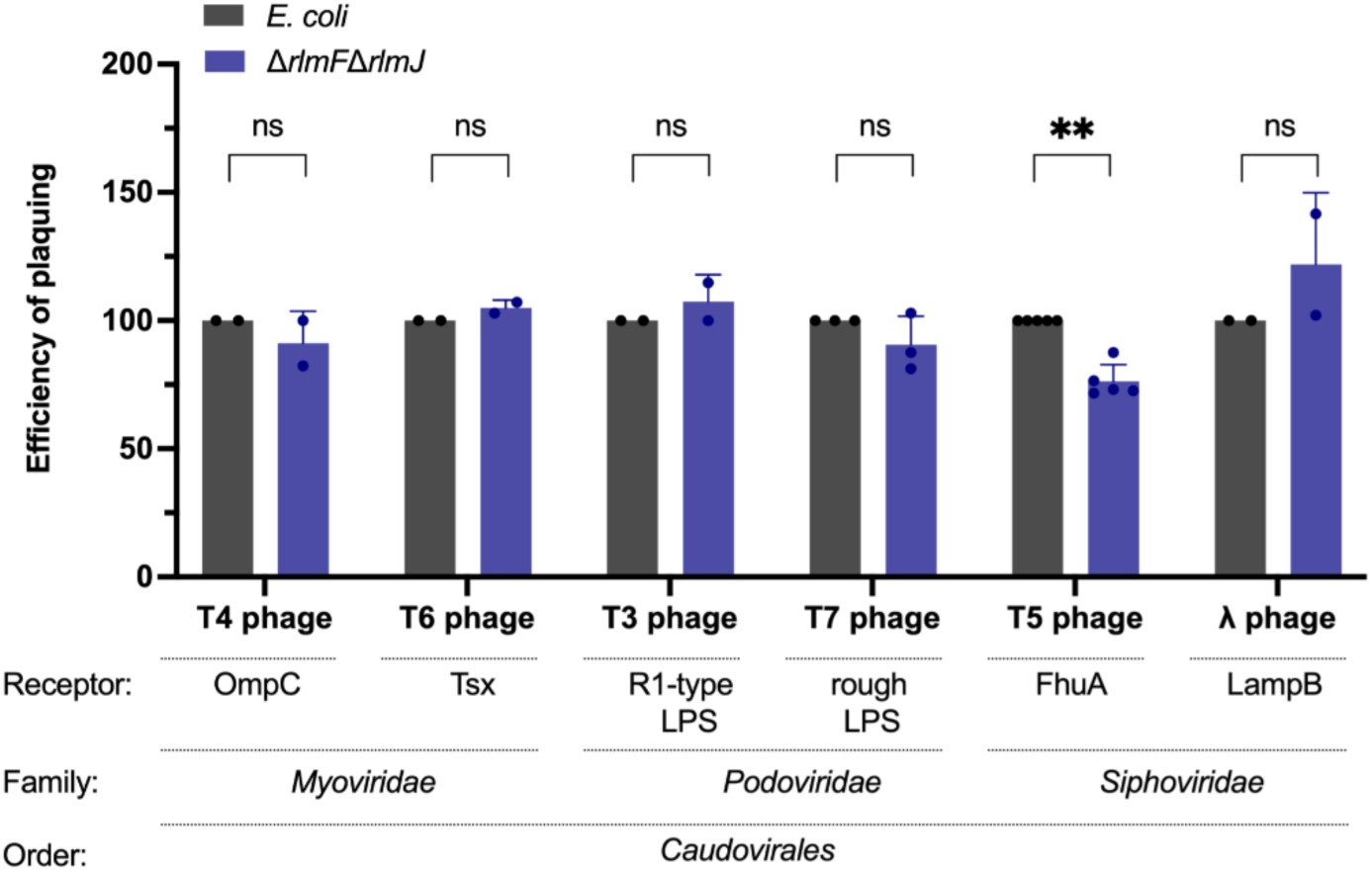
Depletion of m^6^A-RNA in *E. coli* and its effect on infection by different lytic phages. *E. coli* wild type and the Δ*rlmF*Δ*rlmJ* mutant were grown to an OD_600_ = 0.5 before infection with the different phages. The efficiency of plaquing (EOP) was calculated by dividing the number of plaque forming units (PFU) of the Δ*rlmF*Δ*rlmJ* mutant by the number of PFU of the wild type (25). The EOP of the wild type was set to 100. Data are presented as the mean ± standard error of at least two independent experiments. ** *p* < 0.01 (Student’s *t*-test). ns, not significant. The primary receptors, phage families and order are indicated below the diagram.

Of all lytic phages tested, the T5 phage showed significantly reduced plaque formation in the m^6^A-RNA depleted Δ*rlmF*Δ*rlmJ* mutant compared to the wild type, an effect that was not found with the other phages (Fig. 2).

In a more detailed study, we analyzed the infection by T5 phage at different growth stages of *E. coli* (Fig. 3). The number of plaques was always lower in the Δ*rlmF*Δ*rlmJ* mutant compared to the wild type regardless of the growth phase, an effect that was no longer observed in the complemented mutant (ΔΔ-compl.) (Fig. 3). Thus, the lower infection potential of T5 phage is due to the absence of the m^6^A RNA modification.

**FIG 3.**
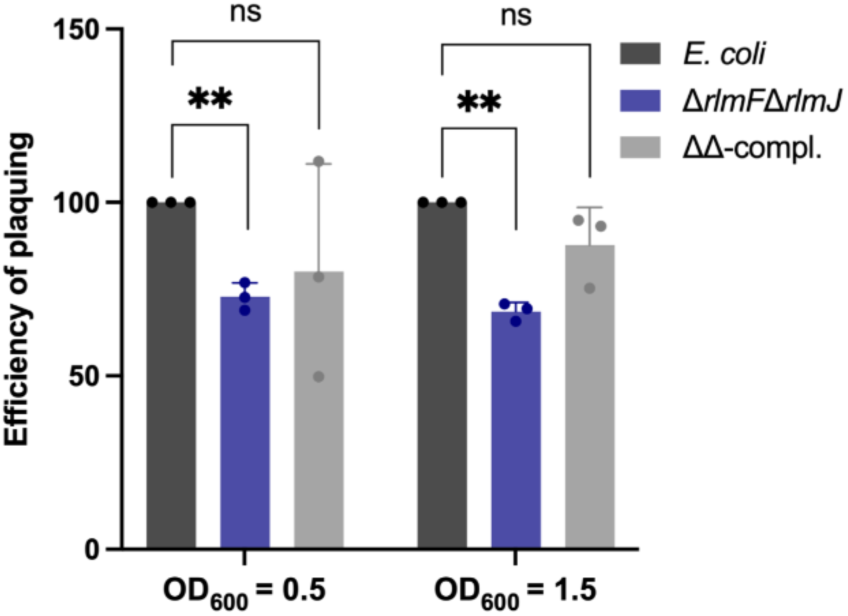
The effect of the growth phase of *E. coli* on T5 phage infection. (A and B) *E. coli* wild type, the Δ*rlmF*Δ*rlmJ* mutant and the complemented mutant (ΔΔ-compl.) were infected in the exponential growth phase (OD_600_ = 0.5) (A) or stationary phase (OD_600_ = 1.5) (B). The efficiency of plaquing was calculated as in Fig. 2. Data are presented as the mean ± standard deviation of three independent experiments. ** *p* < 0.01 (Student’s *t*-test). ns, not significant.

### m^6^A-RNA depletion increases survival of *E. coli* after bacteriophage T5 infection

The reduced efficiency of plaquing of bacteriophage T5 found for the Δ*rlmF*Δ*rlmJ* mutant compared to wild-type *E. coli* (Fig. 2 and 3), prompted us to monitor the survival of both strains at different multiplicities of infection (MOI), which represent the ratio of T5 bacteriophage particles to bacterial cells. In the control experiment (Fig. 4A), bacterial cells proliferated and no lysis was observed. However, the addition of T5 phage resulted in lysis of bacterial cells (Fig. 4B to E), confirming that the observed phenomenon was a consequence of phage infection. Infection of the exponentially grown *E. coli* wild type and the m^6^A-RNA depleted mutant with T5 phage at an MOI of 10 or 1 led to the complete collapse of both populations, with earlier collapse at an MOI of 10 (Fig. 4B and C). In both strains, the excess of T5 phage led to rapid cell death. Similarly, infection at an MOI of 0.1 resulted in earlier collapse of both populations than at an MOI of 0.01 (Fig. 4D and E). Strain-specific differences during infection with T5 phage became evident at an MOI of 0.01 (Fig. 4E). The m^6^A-RNA depleted mutant (Δ*rlmF*Δ*rlmJ*) survived approximately 3 hours longer than the wild type before dramatic cell lysis occurred (Fig. 4E). Furthermore, the cells of the mutant population survived the infection better than the wild type, as about twice as many cells were not lysed (Fig. 4E).

**FIG 4.**
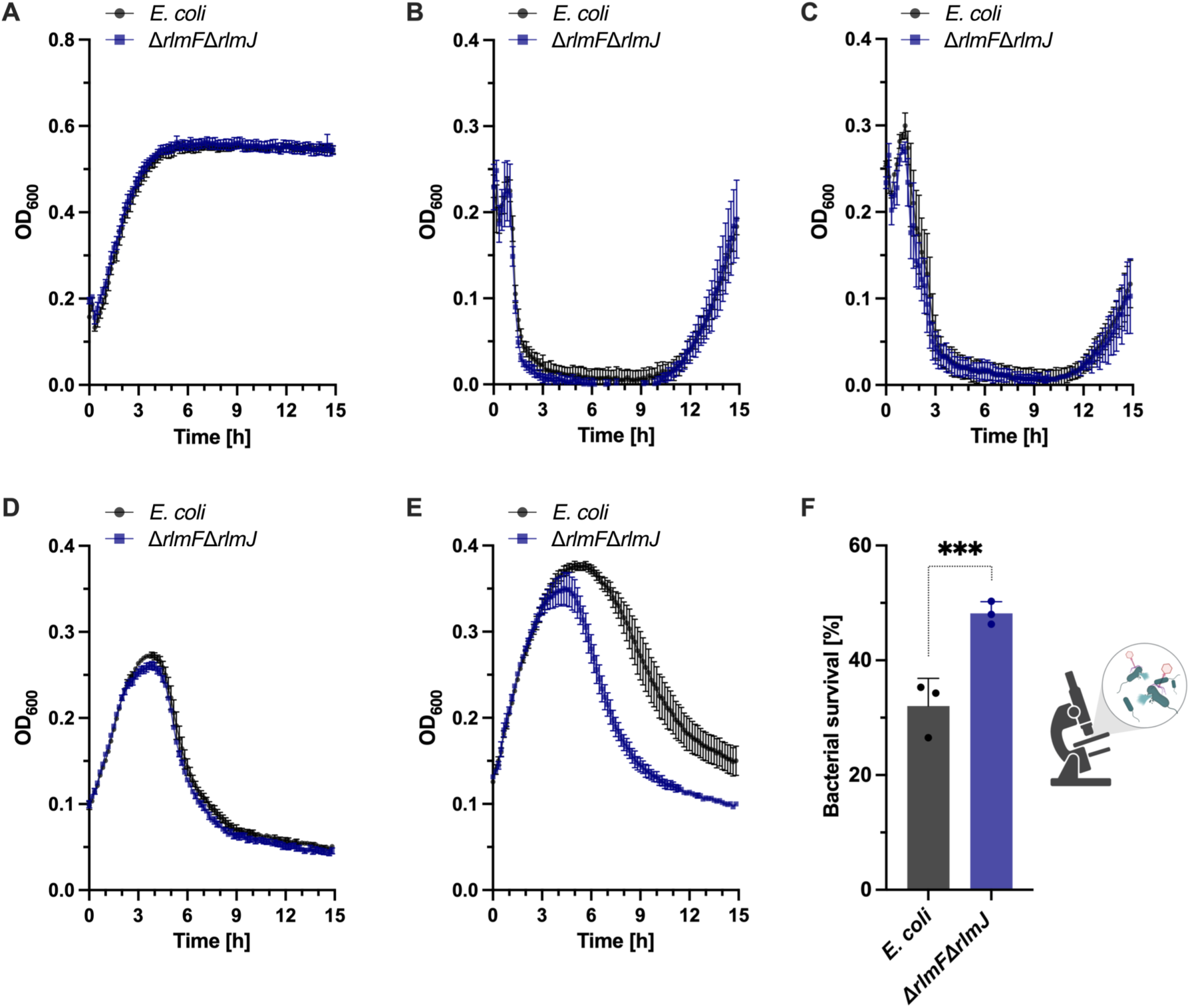
Survival of *E. coli* wild type and Δ*rlmF*Δ*rlmJ* mutant after bacteriophage T5 infection at different MOIs. (A-E) *E. coli* wild type and Δ*rlmF*Δ*rlmJ* mutant were grown to exponential growth phase in Nutrient broth. About 2*10^8^ cells (CFUs) were noninfected (A) or infected with T5 phage at MOI of 10 (B), 1 (C), 0.1 (D) and 0.01 (E). The mixtures were transferred to 96-well plates (150 µL per well) and the OD_600_ was measured every 10 min at 37 °C with continuous shaking. (F) *E. coli* strains were infected with T5 phage at MOI of 10. Single cells were monitored by phase contrast time-lapse microscopy, imaged every 2 min, and the percentage of surviving cells after 2.5 hours was determined. A total of 2,080 wild-type and 1,923 Δ*rlmF*Δ*rlmJ E. coli* cells were quantified at time point 2.5 h post-infection. Data are presented as the mean ± standard error (A-D) or as mean ± standard deviation of the mean (F) of three independent biological replicates.

In addition, we analyzed the time-dependent lysis of the bacteria after T5 infection using phase-contrast time-lapse microscopy. Exponential phase *E. coli* wild type and Δ*rlmF*Δ*rlmJ* mutant were infected with T5 phage at an MOI of 10 to ensure synchronous phage infection or with nutrient broth as a control. After adsorption for 10 min, samples were spotted on an agarose pad and imaged up to 3 hours post-infection. A significantly higher proportion of cells from the /-*rlmF*/-*rlmJ* population survived after 2.5 hours of T5 infection compared to the *E. coli* wild-type population (Fig. 4F). This result is consistent with the survival experiments (Fig. 4E).

### m^6^A-RNA depletion in *E. coli* does not affect T5 phage absorption but diminishes its infection potential

To further investigate T5 phage-host interactions, we investigated adsorption of T5 to *E. coli* wild type and Δ*rlmF*Δ*rlmJ* mutant (Fig. 5A). We observed similar adsorption rates of T5 phage to both host strains, with at least 60 % adsorption within 15 min and 85 % adsorption within 25 min (Fig. 5A). These results suggest that the higher percentage of surviving cells observed in the m^6^A-RNA depleted mutant after T5 phage infection is likely not due to differences in binding to host cells, but is instead related to processes that occur after virus-receptor interaction.

**FIG 5.**
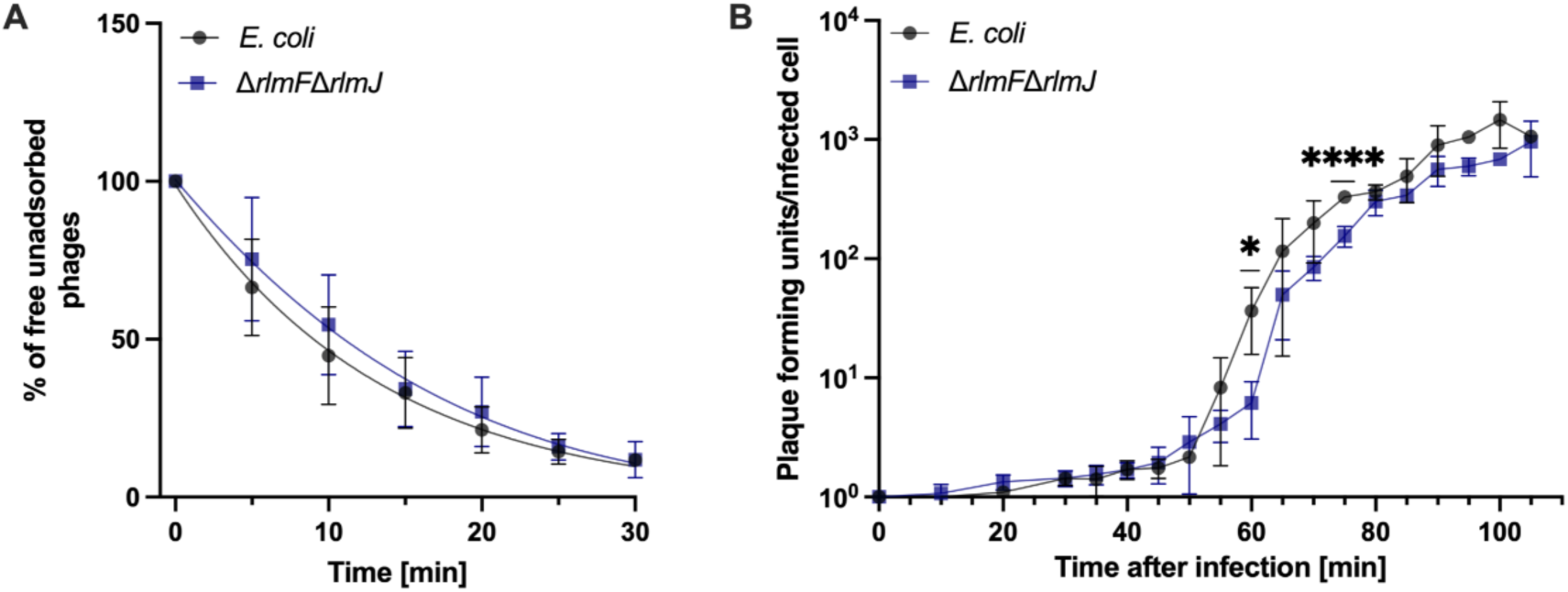
Characterization of T5 phage infection in *E. coli* wild type and Δ*rlmF*Δ*rlmJ* mutant. (A) Adsorption kinetics of T5 phage to wild type and m^6^A-RNA depleted mutant *E. coli* at MOI 0.1. Free phages were collected at the indicated time points after infection (0 min), and the concentration of free phages was determined using plaque assays. (B) One-step growth curve of T5 phage determined for wild type and m^6^A-RNA depleted mutant at MOI 0.01. Data are presented as the mean ± standard deviation of at least three independent experiments * *p* < 0.05; ** *p* < 0.01; **** *p* < 0.0001 (paired Student’s t-test).

To test this, one-step growth curves were performed with the *E. coli* wild type and the Δ*rlmF*Δ*rlmJ* mutant, with the aim to characterize the dynamics of T5 phage replication. In particular, this analysis allowed us to determine (i) the latent period, the time for production of phage components and virion assembly until phage-induced lysis of infected cells, (ii) the rise period, which includes the rise in phage concentration and asynchronous bursts, and (iii) the burst size, a measure of phage virions produced per infected cell (Fig. 5B). For both strains, the latent period was observed to be 45 min, followed by a rise period for another 45 min, followed by a burst period (Fig. 5B). In the Δ*rlmF*Δ*rlmJ* mutant, a significant delay in the increase of the phage titer was observed during the rise period compared to the wild type, which was particularly evident at the 60- and 75-minute time points. This period indicates the rise in phage number due to the release of progeny phage particles by lysis of infected bacterial cells. In addition, the burst size for the *E. coli* wild type was 364 phages compared to 339 phages for the Δ*rlmF*Δ*rlmJ* mutant, a reduction of approximately 7 % per infection cycle. These results suggest that the m^6^A-RNA depleted mutant releases progressively fewer phages over multiple rounds of infection, explaining the higher survival rates (Fig. 4E and F) and the lower number of PFU in the mutant (Fig. 3).

## Discussion

In eukaryotes, m^6^A-RNA modifications are crucial for viral infection and host innate antiviral immunity (30). The effect of this modification is highly dependent on the particular virus and cell types involved, leading to either enhancement or inhibition of viral replication in eukaryotes (30).

To date, there are only a few examples of RNA related modifications known that are important in the context of phage replication (31). Adenosine diphosphate (ADP)-ribosyltransferases (ARTs) of the T4 phage post-translationally modify host proteins with ADP-ribose from the substrate NAD (32, 33). A recent study reported that one of these enzymes, ModB, accepts not only NAD^+^ but also NAD^+^-capped RNA as a substrate (34). Thereby, RNA chains are covalently attached to host proteins in a process, called RNAylation. T4 phages with inactive ModB are characterized by delayed lysis and a reduced number of progenies. The first internal mRNA modification has been described by the group of Michael Laub. They found that the ART CmdT modifies the *N*^6^ position of adenine in GA dinucleotides within single-stranded RNAs leading to an arrest of mRNA translation and inhibition of T4 phage replication (11).

Here, we found that plaquing efficiency of the T5 phage was significantly reduced in a m^6^A-RNA depleted *E. coli* mutant compared to the wild type (Fig. 2 and 3). Consequently, more cells of the mutant survived infection with the T5 phage (Fig. 4). Similarly, the lack of the m^6^A reader/writer YTHDF2 and the m^6^A writer METTL3 in eukaryotic BSC40 cell lines attenuates the replication of dsDNA simian vacuolating virus 40 (SV40) (35).

Since T5 phage efficiently adsorbed to both the wild type and m^6^A-RNA depleted mutant of *E. coli* (Fig. 5A), the greater bacterial survival of the mutant likely resulted from changes during the steps after virus receptor interaction. The one-step growth curve showed a 5-min delay in the rise period of the m^6^A-RNA depleted mutant, which was particularly significant at the 60- and 75-min time points (Fig. 5B). This slight delay in the rise period during T5 phage infection may indicate a delay in phage assembly and/or host cell lysis. Strikingly, when analyzing the behavior of single cells, we observed a substantial diversification of the population during T5 phage infection. A larger proportion of m^6^A-RNA depleted cells survived the infection compared to the wild type. In addition, approximately 30% of wild-type cells resisted lysis (Fig. 4F). We hypothesize that the degree of m^6^A-RNA modification of *E. coli* is heterogeneous and fine-tunes the infection of individual cells by the T5 phage.

In this study, it was shown for the first time that m^6^A-RNA modification of *E. coli* fine-tunes the replication of T5 bacteriophage. Host cells with lower m^6^A levels have a higher chance of surviving phage infection. We hypothesize that a heterogeneous m^6^A modification level in individual cells of the *E. coli* population provides a survival advantage because a subset of the population with lower m^6^A modification levels can escape phage infection and lysis.

## Materials and methods

### Strains, plasmids and oligonucleotides

The bacterial strains, bacteriophage strains and plasmids used in this study are listed in Table 1. Oligonucleotide sequences are listed in Table 2.

**Table 1:**
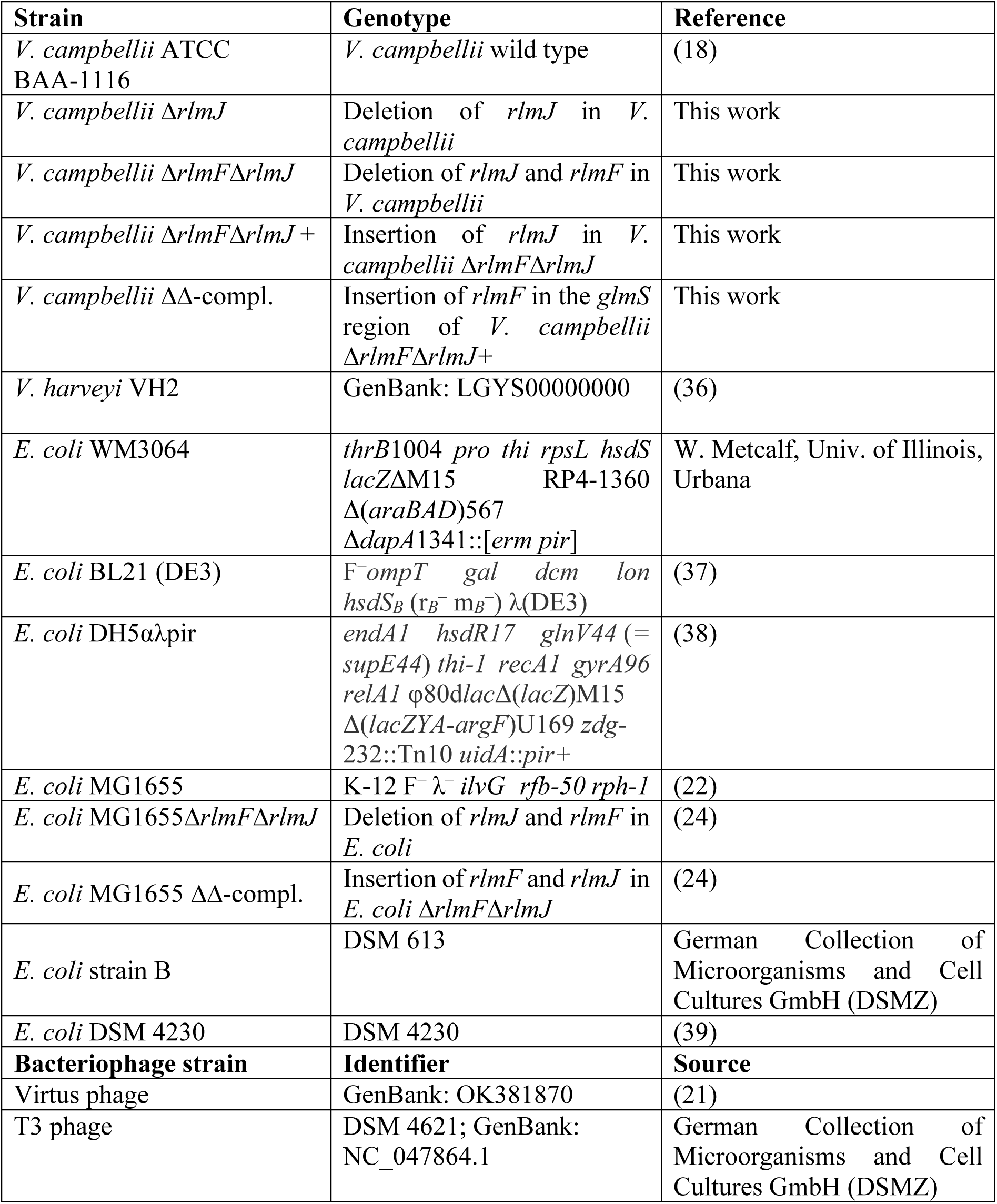
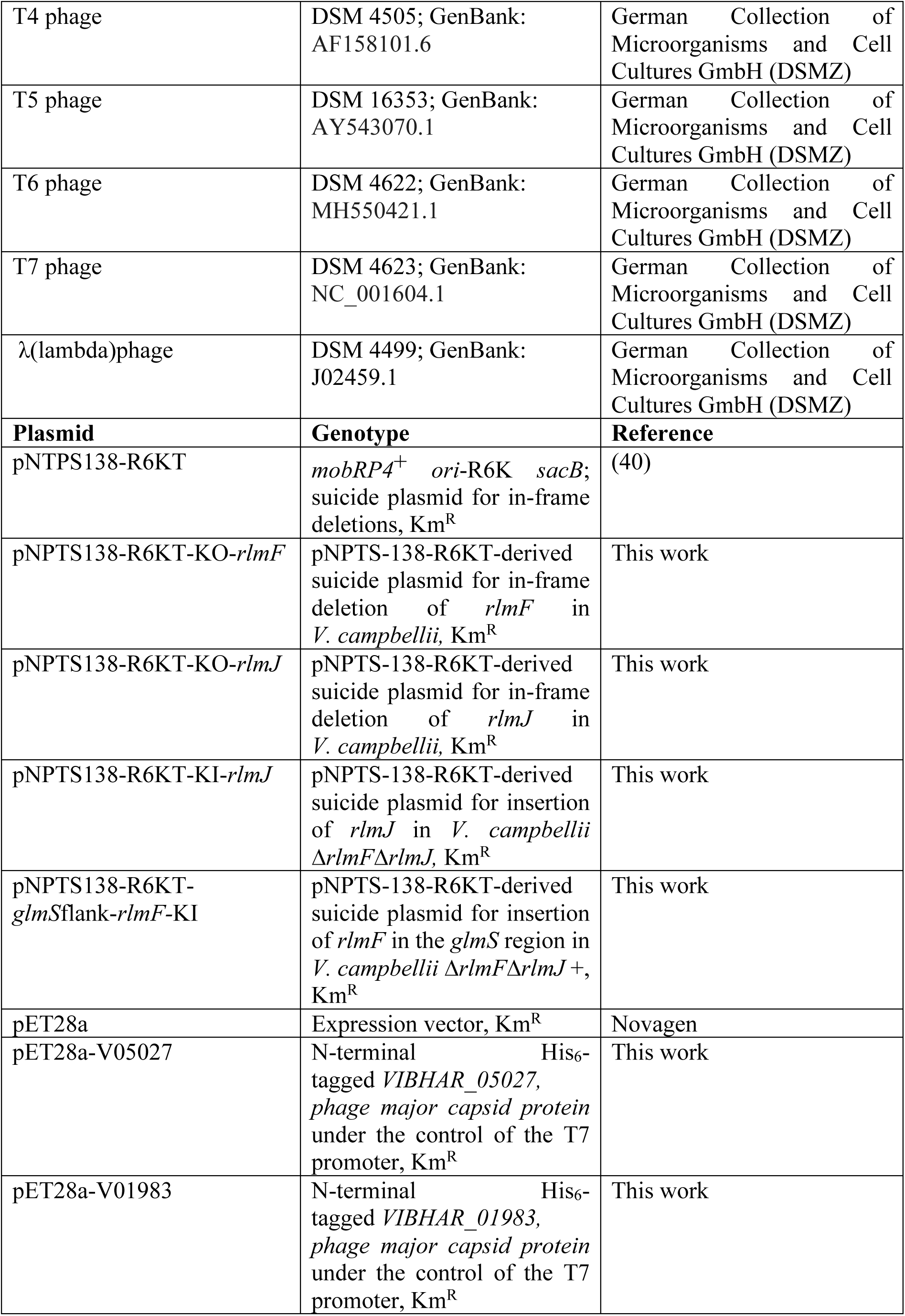
Bacterial strains, bacteriophages and plasmids used in this study.

**Table 2:**
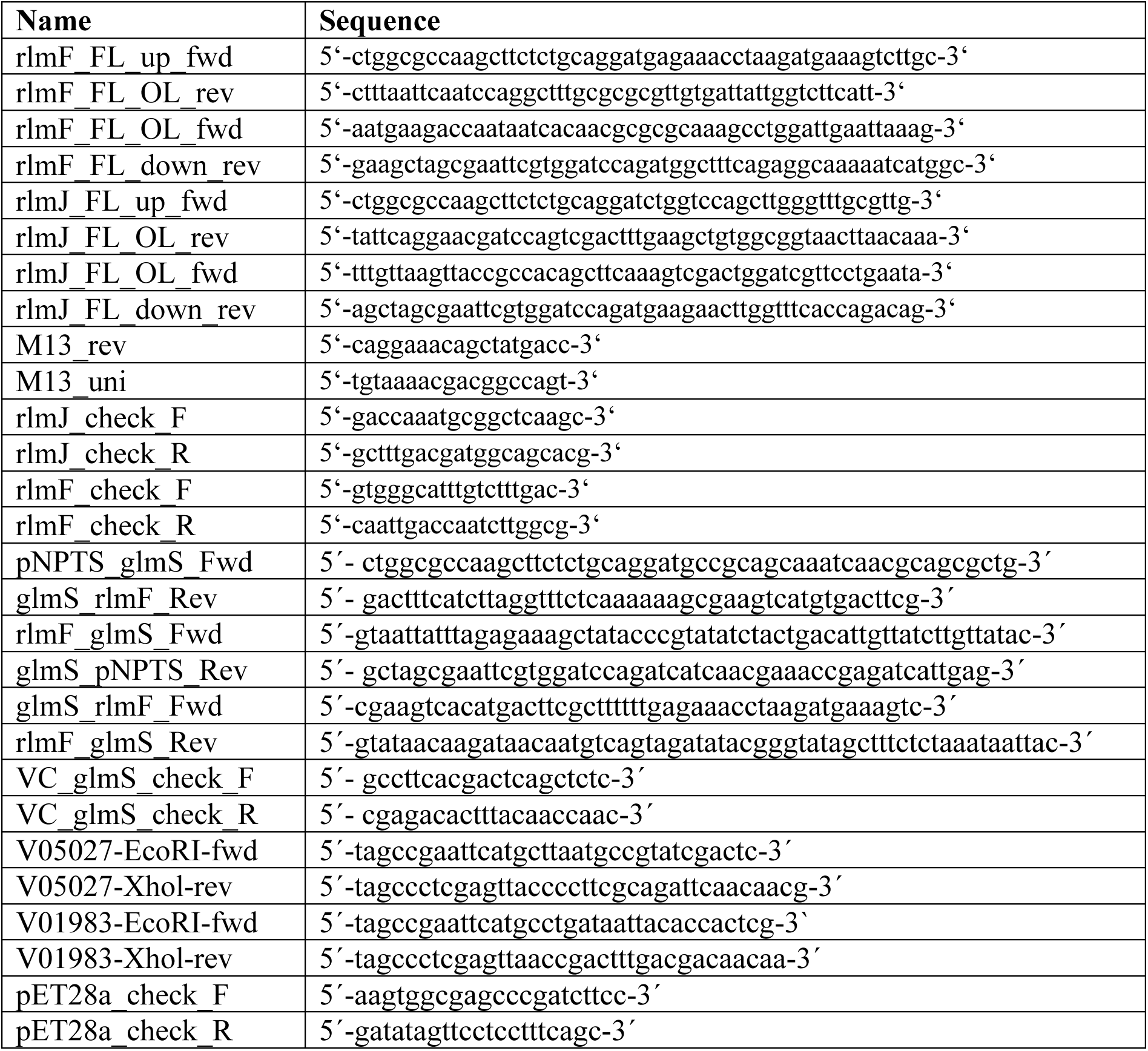
Primers used in this study.

### Plasmid and bacterial strain construction

Molecular methods were carried out according to the manufacturers’ instructions. Chromosomal DNA isolation, plasmid isolation and PCR product clean-up kits were purchased from Süd-Laborbedarf Gauting (SLG) (Gauting, Germany). Enzymes and HiFi DNA Assembly Master Mix were purchased from New England BioLabs (Ipswich, MA, USA).

Construction of the marker-less double deletion mutant *V. campbellii* Δ*rlmF*Δ*rlmJ* was achieved using the suicide plasmids pNPTS138-R6KT-KO-*rlmJ* and pNPTS138-R6KT-KO-*rlmF*. Briefly, 600-bp regions upstream and downstream of *rlmJ* or *rlmF* were amplified by PCR using genomic DNA from *V. campbellii* as a template. The primers rlmJ_FL_up_fwd and rlmJ_FL_OL_rev were used for the upstream fragment of *rlmJ;* rlmJ_FL_OL_fwd and rlmJ_FL_down_rev rev for the downstream fragment of *rlmJ*; rlmF_FL_up_fwd and rlmF_FL_OL_rev for the upstream fragment of *rlmF*; and rlmF_FL_OL_fwd and rlmF_FL_down_rev for the downstream fragment of *rlmF*. After purification, the PCR fragments were cloned by Gibson assembly (41) into the suicide plasmid pNPTS138-R6KT, which was first linearized by digestion with EcoRV. The correct plasmids were verified by colony PCR and sequencing using primers M13_rev and M13_uni.

The suicide plasmid pNPTS138-R6KT-KO-*rlmJ* was next introduced into wild-type *V. campbellii* by conjugative mating using *E. coli* WM3064 as a donor in LB medium containing diaminopimelic acid (DAP) as previously described (42). Briefly, single-crossover integration mutants were selected on LM plates that contained kanamycin but lacked DAP. Single colonies were grown for two days without antibiotics and plated on LB containing 10% (w/v) sucrose to select for plasmid excision. Kanamycin-sensitive colonies were checked for gene deletion by colony PCR using primers bracketing the deletion sites. Deletion of *rlmJ* was verified by colony PCR and sequencing using the primers rlmJ_check_F and rlmJ_check_R. *rlmF* was then deleted through the introduction of the suicide plasmid pNPTS138-R6KT-KO-*rlmF* into *V. campbellii* Δ*rlmJ* via conjugative mating using *E. coli* WM3064 as a donor as described above. Deletion of *rlmF* was verified by colony PCR and sequencing using the primers rlmF_check_F and rlmF_check_R.

Complementation of the *rlmJ* deletion in *V. campbellii* Δ*rlmF*Δ*rlmJ* was achieved with the suicide plasmid pNPTS138-R6KT-KI*-rlmJ*. Briefly, *rlmJ* and 600-bp upstream and downstream flanking regions were amplified by PCR using genomic DNA from *V. campbellii* as a template and the primers rlmJ_FL_up_fwd and rlmJ_FL_down_rev. After purification, the 2030-bp long PCR fragment was cloned by Gibson assembly (41) into the suicide plasmid pNPTS138-R6KT, which was linearized by digestion with EcoRV. The correct plasmids were verified by colony PCR and sequencing using primers M13_rev and M13_uni. Construction of the suicide plasmid pNPTS138-R6KT-*glmS*flank-*rlmF-*KI was slightly different due to the presence of transposable elements in the vicinity of the gene. Briefly, the regions surrounding *glmS* and *rlmF* (600 bp upstream and 24 bp downstream) were amplified to generate the suicide plasmid pNPTS138-R6KT-*glmS*flank-*rlmF* using *V. campbellii* genomic DNA as a template. Primer pairs used for amplification of *glmS* overhang fragments were pNPTS_glmS_Fwd and glmS_rlmF_Rev (upstream of *rlmF*) and rlmF_glmS_Fwd and glmS_pNPTS_Rev (downstream from *rlmF*). The primers glmS_rlmF_Fwd and rlmF_glmS_Rev were used to amplify *rlmF*. These DNA fragments were assembled via Gibson assembly (41) into the pNPTS138-R6KT plasmid linearized with EcoRV. The plasmid was verified by colony PCR and sequencing using primers M13_rev and M13_uni.

To complement Δ*rlmF*Δ*rlmJ*, the suicide plasmid pNPTS138-R6KT-KI*-rlmJ* was introduced into *V. campbellii* Δ*rlmF*Δ*rlmJ* by conjugative mating using *E. coli* WM3064 as a donor in LB medium containing DAP as described above. Insertion of *rlmJ* was verified by colony PCR and sequencing using the primers rlmJ_check_F and rlmJ_check_R. *rlmF* was then added via introduction of suicide plasmid pNPTS138-R6KT-*glmS*flank-*rlmF-*KI to *V. campbellii* Δ*rlmF*Δ*rlmJ+* by conjugative mating with *E. coli* WM3064 as described above. Insertion of *rlmF* in the *glmS* region of *V. campbellii* was verified by colony PCR and sequencing using the primers VC_glmS_check_F and VC_glms_check_R.

To generate plasmids encoding N-terminal 6His-tagged versions of major capsid proteins from *Vibrio* kappa-like prophage and ΦHAP-1-like prophage (*VIBHAR_05027* and *VIBHAR_01983,* respectively), *V. campbellii* genomic DNA was used as a template with primers V05027-EcoRI-fwd and V05027-Xhol-rev (*VIBHAR_05027*) or V01983-EcoRI-fwd and V01983-Xhol-rev (*VIBHAR_01983*). These two genes were cloned into the pET-28a vector using EcoRI and XhoI as restriction sites; the presence of the correct insert was confirmed by colony PCR and sequencing using primers pET28a_check_F and pET28a_check_R.

### Mass spectrometry (MS) analysis of mRNA and rRNA methylation in *V. campbellii*

To identify m^6^A methylation of the rRNA and mRNA moieties in wild type and Δ*rlmF*Δ*rlmJ* mutant *V. campbellii*, total RNA was isolated and processed as previously described (12). In brief, *V. campbellii* cells were grown to an optical density at 600nm (OD_600_) of 1 and harvested by centrifugation. Total RNA was isolated and tRNA was removed by size exclusion chromatography, resulting in a fraction containing mRNA and rRNA moieties. This RNA fraction was hydrolyzed and quantitatively analyzed by liquid chromatography-tandem MS (LC-MS/MS) (12).

### Overproduction and purification of recombinant phage capsid proteins

*E. coli* BL21 (DE 3) was transformed with the vectors pET28a-V05027 and pET28a-V01983 to overexpress the major capsid proteins of *Vibrio* kappa-like prophage and ΦHAP-1-like prophage, respectively. Cells were cultivated in LB supplemented with kanamycin (50 mg/mL) at 37 °C to an OD_600_ of 0.5. Isopropyl-β-D-1-thiogalactopyranoside (IPTG) (0.5 mM) was added to the culture to induce *VIBHAR_05027* or *VIBHAR_01983* expression at 37 °C for 2.5 h. Cells were harvested (5000 × *g*, 20 min, 4 °C), resuspended, and disrupted in ice-cold disruption buffer (50 mM Tris-HCl [pH 7.5], 10% (v/v) glycerol, 10 mM magnesium chloride, 1 mM dithiothreitol, 0.5 mM phenazine methosulfate [PMSF], 3 mg DNase and 100 mM NaCl) using a high-pressure cell disrupter (Constant Systems Limited, Daventry, UK). Cell debris and intact cells were removed by centrifugation (5000 × *g*, 20 min, 4 °C) and the pellet was solubilized in 6 M urea and 10 mM Tris-HCl (pH 7.5) overnight with shaking at 4 °C. Any precipitates or unsolubilized aggregates were removed by ultracentrifugation (20,000 × *g*, 15 min, 4 °C) and the supernatant was loaded onto a Ni-nitrilotriacetic acid (NTA) column (Qiagen, Hilden, Germany). After a washing step (6 M urea, 10 mM Tris-HCl [pH 7.5] and 40 mM imidazole), the recombinant protein was eluted with elution buffer (6 M urea, 10 mM Tris-HCl [pH 7.5] and 200 mM imidazole).

### Generation of polyclonal rabbit antibodies against recombinant phage capsid proteins

Customized polyclonal rabbit antibodies against 6His-VIBHAR_05027 and 6His-VIBHAR_01983 were purchased from Kaneka Eurogentec (Seraing, Belgium). Heterologously produced and purified 6His-VIBHAR_05027 or 6His-VIBHAR_01983 were used as antigens in a Speedy 28-day Immunization Program with two rabbits per antigen as hosts. The specificity of the polyclonal antibodies against 6His-VIBHAR_05027 and 6His-VIBHAR_01983 was verified by western blot analysis as described below.

### Sodium dodecyl sulphate-polyacrylamide gel electrophoresis (SDS-PAGE) and western blot analysis of 6His-VIBHAR_05027 and 6His-VIBHAR_01983

#### SDS-PAGE (43)

Stacking gels consisted of 4% (w/v) acrylamide (in 50 mM Tris [pH 6.8]) and resolving gels of 12.5% (w/v) acrylamide (in 300 mM Tris [pH 8.8]) and were run in a Tris-glycine buffer (25 mM Tris, 192 mM glycine, 0.1% [w/v] SDS [pH 8.3]). Gels were stained overnight in Coomassie staining solution (0.25% [w/v] Coomassie Brilliant Blue R-250, 9.2% [v/v] concentrated acetic acid and 45.4% [v/v] ethanol) and destained in destaining solution (10% [v/v] acetic acid and 40% [v/v] ethanol).

#### Western blot analysis

Proteins were transferred to a nitrocellulose membrane using the Trans-Blot Turbo Transfer System (Bio-Rad, Hercules, CA, USA) and then blocked with 5% (w/v) milk powder prepared in Tris-buffered saline (TBS) (pH 7.6) containing 0.1% (v/v) Tween-20 (TBST) for 1 h at room temperature. 6His-VIBHAR_05027 and 6His-VIBHAR_01983 were detected by incubating the membrane in TBST with primary polyclonal antibodies (Kaneka Eurogentec, Seraing, Belgium) or primary monoclonal antibodies against the 6His-tag (Thermo Fisher Scientific, Waltham, MA, USA). Membranes were then incubated with the alkaline phosphatase-conjugated goat anti-rabbit IgG (Thermo Fisher Scientific, Waltham, MA, USA) and developed using substrate solution (50 mM sodium carbonate buffer [pH 9.5], 0.1% [w/v] nitroblue tetrazolium and 5 mg/mL 5-bromo-4-chloro-3-indolyl phosphate). PageRuler Prestained Protein ladder (10-180 kDa) (Thermo Fisher Scientific, Waltham, MA, USA) was used as a size standard ladder.

### Phage induction assays

Wild type, Δ*rlmF*Δ*rlmJ* mutant and complemented mutant (ΔΔ-compl.) *V. campbellii* were exposed to stress conditions to assess the effect of m^6^A RNA modification on prophage induction. Bacterial strains grown in LM medium to an OD_600_ of 0.4 were exposed to either oxidative stress (by addition of 1 µg/mL mitomycin C), heat stress (45 °C) or neither (control). After 30 min of oxidative stress, mitomycin C was removed from the sample by centrifugation (5000 × *g*, 10 min, 4 °C) and the pellet was resuspended in fresh, pre-warmed LM medium. After 30 min of heat stress, the samples were shifted to 30 °C. After 2 h of incubation under physiological conditions, the supernatant containing the phage particles was collected (5000 × *g*,15 min, 4 °C), passed through a 0.22 µm Millipore Steriflip Vaccum filter (MilliporeSigma, Burlington, MA, USA) and stored at 4 °C.

To follow cell lysis caused by the release of lysogenic phages, wild type, Δ*rlmF*Δ*rlmJ* mutant and complemented mutant (ΔΔ-compl.) *V. campbellii* were exposed to 1 µg/mL mitomycin C after strains were grown in LM medium to an OD_600_ of 0.4. After 30 min of oxidative stress, mitomycin C was removed from the sample by centrifugation (5000 × *g*, 10 min, 4 °C) and the pellet was resuspended in fresh, pre-warmed LM medium. The mixtures were transferred to 96-well plates (150 µL per well) and optical density (OD_600_) was measured every 10 min using a NanoQuant Infinite M200PRO plate reader (Tecan, Männedorf, Switzerland) at 37 °C with continuous shaking.

### Indirect enzyme-linked immunosorbent assay (ELISA)

Indirect ELISA was performed according to described protocols (44, 45) with slight modifications for the detection of the major capsid proteins of *Vibrio* kappa-like and ΦHAP-1-like prophages from *V. campbellii*. In brief, 96-well F-bottom, polystyrene, chimney well, black, medium binding microplates (Greiner Bio-One GmbH, Kremsmünster, Austria) were coated with 100 µL of supernatant (derived from the phage induction assays described above) and 0.05 M carbonate buffer (pH 9.6) and then incubated overnight at 4 °C. After washing three times with TBST, each well of the plate was blocked with 200 µL of 5% (w/v) milk powder prepared in TBST for 1 h at 30 °C. After washing three times with TBST, 100 µL of VIBHAR_05027- or VIBHAR_01983-positive serum in 5% (w/v) milk powder with TBST was added to each well and the plate was incubated for 1 h at 30 °C. After three washes with TBST, 100 µL of alkaline phosphatase-conjugated goat anti-rabbit IgG (Thermo Fisher Scientific, Waltham, MA, USA) diluted in 5% (w/v) milk powder in TBST was added to each well, followed incubation at 30 °C for 1 h and five washes with TBST. Subsequently, 100 µL of the substrate para-nitrophenyl phosphate (PNPP) (Thermo Fisher Scientific, Waltham, MA, USA) was added to each well and incubated for 30 min in the dark. Last, 50 µL of 2 N sodium hydroxide was added to each well to stop the reaction. The absorbance was immediately measured at 405 nm in a NanoQuant Infinite M200PRO plate reader (Tecan, Mäannedorf, Switzerland). All samples were analyzed in technical duplicates.

Standard curves generated with purified major capsid proteins were included in each ELISA run to estimate the concentration of the two major capsid proteins in the sample.

### Propagation and determination of phage titer using plaque assay

To obtain high-titer phage stocks, propagation of all phages mentioned in this work was performed using the plate lysate method as described (46). The propagation host used for Virtus phage was *V. harveyi* VH2; the λ phage host was *E. coli* DSM 4230; and the host for all T-phages was *E. coli* B strain (DSM 613).

To determine the phage titer, a double agar overlay plaque assay was performed as described (47). In brief, 100 µL of exponential-phase bacterial culture was mixed with 100 µL of diluted phages in soft agar, which was then poured onto a solid medium. The plates were incubated overnight and the number of plaques formed was counted to determine the titer.

### Efficiency of plaquing (EOP)

EOP was determined after performing the plaque assay described above, using phage lysates to infect wild type and Δ*rlmF*Δ*rlmJ* mutants of *V. campbellii* and *E. coli*. The EOP was calculated by dividing the number of plaque forming units (PFU) of the Δ*rlmF*Δ*rlmJ* mutant by the number of PFU of the wild type (25). The EOP of the wild type was set to 100.

### Bacterial survival after T5 phage infection

*E. coli* wild type and Δ*rlmF*Δ*rlmJ* mutant were grown to exponential growth phase in Nutrient broth (N7519, Sigma-Aldrich, St. Louis, MO, USA). Cells (about 2*10^8^ colony forming units [CFU]/mL) were infected with T5 phage at a multiplicity of infection (MOI) of 10, 1, 0.1 or 0.01 to monitor host dynamics. Mixtures were transferred to 96-well plates (150 µL per well) and optical density (OD_600_) was measured every 10 min using a NanoQuant Infinite M200PRO plate reader (Tecan, Männedorf, Switzerland) at 37 °C with continuous shaking.

### Synchronized infection assay

The synchronized infection assay was performed as previously described (48, 49). *E. coli* wild type and Δ*rlmF*Δ*rlmJ* mutant were grown as described above and infected with T5 phage at a MOI of 10. Samples were collected at 0 min and 5 min after infection and immediately plated for CFU enumeration. Three biologically independent infection experiments showing a CFU reduction of at least 95% within 5 min of infection were considered as synchronized infection.

### Time-lapse phase contrast microscopy

Phase-contrast time-lapse microscopy was used to follow T5 phage-mediated bacterial lysis using a modified protocol described by Mandal and colleagues (50). *E. coli* wild type and Δ*rlmF*Δ*rlmJ* mutant were grown as described above and infected with T5 phage at a MOI of 10 (or with an equal volume of phage diluent as a control). After a 10 min incubation for initial adsorption at 37 °C under agitation, 2 µL of cells were spotted on pads composed of Nutrient broth (N7519, Sigma-Aldrich, St. Louis, MO, USA) and 1% (w/v) agarose. These pads were placed on microscopic slides and covered with a coverslip. Within 20 minutes, images were taken on a Leica DMi8 inverted microscope equipped with a Leica DFC365 FX camera (Wetzlar, Germany). Microscopic phase-contrast images were captured every two min in several positions up to 3 h with a constant temperature of 37 °C using an incubator (PeCon, Erbach, Germany) around the DMi8 microscope. To set the focus plane, automatic autofocus was performed at every position and every time point using the adaptive focus control (AFC) and closed-loop focus system of the DMi8 microscope. To quantify non-lysed cells, phase-contrast images were analyzed using the plugin MicrobeJ (51) in ImageJ (52). The default MicrobeJ settings were used for cell segmentation (Fit shape, rod-shaped bacteria) apart from the following settings: area: 1-max µm; length: 2.5–20 µm; width: 0.4–2 µm; curvature 0.0– 0.15, angularity 0.0–0.25 and intensity 0.0–2000.

### T5 bacteriophage adsorption assay

Adsorption assays were performed as previously described (53). *E. coli* wild type and Δ*rlmF*Δ*rlmJ* mutant were grown as described above and infected with T5 phage at an MOI of 0.1. Free phages were sampled immediately post-infection (0 min), then every 5 min for 30 min using 0.22-µm sterile filtration. Plaque assays (47) were performed as described above to enumerate the free phage concentration.

### One-step growth curve

One-step growth curves were performed as described previously (48, 54). *E. coli* wild type and Δ*rlmF*Δ*rlmJ* mutant were grown as described above, infected with T5 phage at an MOI of 0.001 and incubated at 37 °C for 10 min. After initial adsorption, the remaining free phages were removed via centrifugation (5000 × *g*, 5 min, 4 °C) and the pellet was resuspended in fresh pre-warmed Nutrient broth. Samples were incubated at 37 °C with shaking and collected (including t = 0) at 10-min intervals over 30 min, followed by 5-min intervals up to 105 min. Plaque assays (47) were performed as described above immediately after sample collection.

### Statistical analysis and data presentation

All numerical data were analyzed and graphs were prepared with GraphPad Prism version 10.0.2 (GraphPad Software, La Jolla, CA, USA). Statistics were performed using paired and unpaired Student’s *t-*test. Microscopy images were prepared from a total of 2,080 *E. coli* wild type and 1,923 Δ*rlmF*Δ*rlmJ* mutant cells and analyzed with MicrobeJ. All figures were generated using Affinity Designer version 2.2.1 (Serif, West Bridgford, UK).

All experiments were repeated at least three times to ensure reproducibility.

## Acknowledgements

We thank Korinna Burdack and Inès Wullkopf for conducting plaque assays, Dimitar Petrov for isolating RNA, and Siobhan Cusack for proofreading the manuscript. B.S. acknowledges the support of the Graduate School Life Science Munich (LSM).

This work was financially supported by the Deutsche Forschungsgemeinschaft (DFG, German Research Foundation): Project numbers 464582101 (JU270/21-1; BR6622/1-1) and 325871075 (SFB 1309).

B.S., S.B. and K.J. conceptualized the study. B.S., S.B., T.C., S.R. and K.J. designed the methodology. B.S. performed the investigation. B.S., S.B. and K.J. wrote the original draft, B.S., S.B., T.C., S.R. and K.J. reviewed and edited the manuscript. K.J. and S.B. acquired funding, provided resources, and supervised the study.

## Data availability

Additional information required to reanalyze the data reported in this paper is available from the lead contacts upon request.

